# Developing a model of temporomandibular disorder in the common marmoset using nerve growth factor

**DOI:** 10.1101/2025.08.20.671273

**Authors:** Erin J. Holzscherer, Rhonda Kersten, Mathilde Bertrand, Jibran Y. Khokhar, Brian E. Cairns, J. Andrew Pruszynski, David A. Seminowicz

## Abstract

Developing an animal model that more closely represents the human multidimensional pain experience is an important step towards addressing the current chronic pain crisis. The common marmoset has potential as this model species given its biological, neurological and phylogenetic similarity to humans. Here, we developed a model of myofascial temporomandibular disorder (TMD) in the marmoset by injecting nerve growth factor (NGF) into the superficial masseter. Following injection, animals showed reduced mechanical withdrawal thresholds at 5 μg and 10 μg doses of NGF and changes in circadian rhythm and feeding initiation following injection of 10 μg of NGF. Animals did not show evidence of jaw dysfunction, masticatory alterations, or grimace during novel behavioural assays. The model is transient, with pain resolution occurring approximately 7 days after onset, which allows for repeated testing on the same animal. This same NGF-TMD model has been previously validated in rodents and humans and presents an opportunity for forward and reverse translation to examine mechanisms, develop relevant pain assessment tools, and ultimately test novel treatments for TMD and other musculoskeletal pain conditions.

**New & Noteworthy:** We developed a long-lasting but transient (∼7 days) model of myofascial temporomandibular disorder pain in marmosets. Mechanical hypersensitivity and changes to circadian activity and spontaneous eating behaviours were observed. There was no evidence of jaw dysfunction, altered food preference or changes in grimace. The NGF-TMD model can be translated to the marmoset with the potential for investigating mechanisms and novel interventions for TMD.

## Introduction

Chronic pain is associated with substantial personal and societal burden, and there is a recognized need for novel relevant and efficacious interventions (1,2). Many interventions continue to fail in clinical trials partially due to the lack of predictive animal models (3–7). While animal models provide a necessary avenue to assess interventions prior to clinical application, many current rodent models do not address the multidimensional cognitive and affective components of the human pain experience (1,3–6,8). Specifically, while rodent models demonstrate mechanical hypersensitivity similar to human pain patients, the spontaneous, behavioural aspects of pain are difficult to reproduce in rodents (9). Recent studies have recognized the need for analyzing spontaneous or affective pain behaviours in rodent models by investigating grimace scales (8,10), masticatory alterations evoked by orofacial pain (11,12), quality-of-life factors (9,13,14) and operant behavioral assays (6,10,15,16). However, many of these studies show inconsistent findings and continue to fail as forward translational models. Inherently, the biological and neurological dissimilarities between rodents and humans cannot be alleviated by changing behavioural measures; treatments for spontaneous rodent behaviour may not address the affective or neurological impairments experienced by human patients. This discrepancy can be addressed by exploring other species, such as non-human primates (NHPs), that share more neurological and behavioral similarities to humans.

The common marmoset (*Callithrix jacchus*) has the potential to represent a holistic pain model considering their neurological, behavioural and phylogenetic similarities to humans (17). Their small size and high fecundity give them an advantage over traditional Old World primates (17,18) while conserving many primate-specific anatomical and functional brain network properties (19,20). Additionally, marmosets can be trained on behavioural assays including reaching tasks (21–25), touch screen tasks (25–27), as well as more complex cognitive tasks (28–30). These features demonstrate the potential for a marmoset model to address possible behavioural, cognitive, and mechanistic aspects of the pain experience. However, developing a marmoset pain model relies on available species-specific pain assessment tools which are currently lacking. Systematic approaches to measuring pain in the marmoset are almost non-existent (31,32) and marmoset pain assessment relies heavily on subjective observations such as reduced alertness or withdrawal, although these measures are not specific to pain (i.e. social isolation, stress) (31,33). Therefore, it is important to establish new and objective measures of pain assessment in the marmoset to ensure pain can be recognized quickly and accurately for improved welfare and developing an efficacious chronic pain model.

Given the recognized need for both a relevant animal pain model and accurate marmoset pain assessment tools, the goal of this study was to pilot a validated pain model of temporomandibular disorder (TMD) in the common marmoset using a biologically relevant pain stimulant. In Experiment 1, we injected different concentrations of nerve growth factor (NGF) into the superficial masseter muscle to establish an effective dose and measured subsequent eating behaviours using novel in-lab behavioural assays. In Experiment 2, we used the dose of NGF established from Experiment 1 to assess naturalistic home cage behaviours. The action of NGF has been shown to cause mechanical hyperalgesia, allodynia, jaw dysfunction, masticatory alterations and changes in eating behaviour in rodents (34–36) and humans (37–40) As such, our approach is intended to create a back- and forward- translation model by using a validated pain stimulant to develop relevant behavioural assays and explore naturalistic TMD-related pain behaviours in the marmoset.

## Methods

### Animals

The NGF-TMD model was developed with two adult marmosets (1 female, 7 years, 460 g; 1 male, 4 years, 388 g). Animals were pair housed in a temperature (25 ± 1° C) and humidity (55 ± 15%) controlled facility with a 12 h light-dark cycle. Animals had unrestricted access to food and water. All procedures followed protocols approved by the Animal Care Committee of The University of Western Ontario (Protocol # 2023-065) in accordance with the Canadian Council on Animal Care policy.

#### Experiment 1: Dose effects and in lab behavioural analysis

##### NGF administration

Nerve growth factor (NGF) (0.1 g; human, MilliporeSigma) was reconstituted with 1 ml of sterile 1x phosphate buffered saline (PBS, 100 μl/ml), aliquoted (10 μl vial), and stored at -20°C for later use. On Day 1 and Day 3 of experimental conditions, animals were anesthetized using Alfaxalone (3-4 mg/kg), Medetomidine (0.03 mg/kg), and maintained on Isoflurane (1%). Heart rate, blood oxygen levels and temperature were monitored throughout the procedure. Once fully anesthetized, 100 μl of NGF (25 μg/mL, 50 μg/mL or 100 μg/mL) or 100 μl of PBS was injected into the left or right masseter. The muscle was located using landmarks established during dissections on animals euthanized for other studies and this dose was established by injecting animals approximately every 4 weeks with incremental increases to concentration and volume (Supplementary Table 1) until a quantifiable response was observed. Following NGF injections, animals were given atipamezole hydrochloride (Revertor; volume/volume of medetomidine; Modern Veterinary Therapeutics, LLC) and monitored until fully awake. Once awake, animals were placed back in the home cage to fully recover. The experimenter was blinded to the side of NGF injection during the procedure and duration of testing.

##### Behavioural testing

Animals received extensive habituation using positive reinforcement training prior to any injection of NGF and all behavioural assays were developed specifically for this model using weeks of trial-and-error testing. During both baseline and treatment conditions, animals received daily behavioural testing in the lab, including the days of injection (Day 1, Day 3) until all measures returned to baseline as described below (Figure 1A). All sessions were video recorded and analyzed following testing. On injection days, testing was done when the animal was fully recovered from anesthesia.

**Figure 1.**
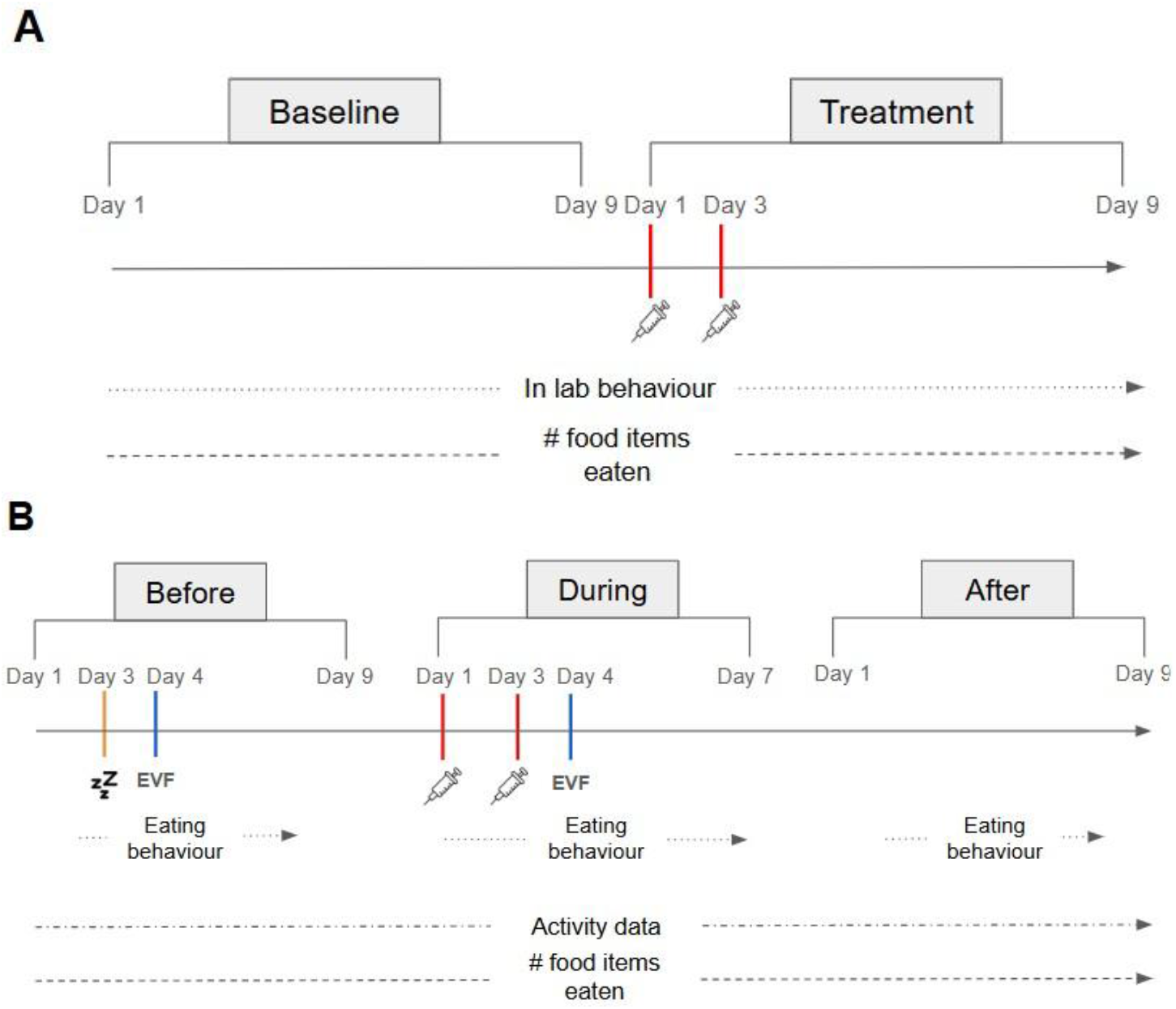
Experimental timelines. (A) Experiment 1 and (B) Experiment 2.

###### Mechanical withdrawal test

Mechanical withdrawal was evaluated using an electronic von Frey (EVF, Bioseb) While seated in the primate chair, animals were provided a constant rate of acacia gum (0.8 ml/min) to maintain head stillness. Once engaged with the reward, the EVF was manually applied with increasing pressure to the masseter muscle (Figure 2A). The pressure at which the animal withdrew their head (withdrawal) was automatically recorded. This was repeated 6 times for each (left, right) muscle with a minimum of 30 seconds between each withdrawal. If an external stimulus (i.e. noise outside the lab) provoked a withdrawal, this was considered a “false withdrawal” and was excluded and repeated. Mechanical withdrawals were normalized to an average baseline threshold taken from nine days of pre-NGF behavioural testing to determine the change in withdrawal thresholds for each side. Relative withdrawal thresholds can be used as a measure of quantitative sensory testing in conditions of unilateral affliction (41).

**Figure 2.**
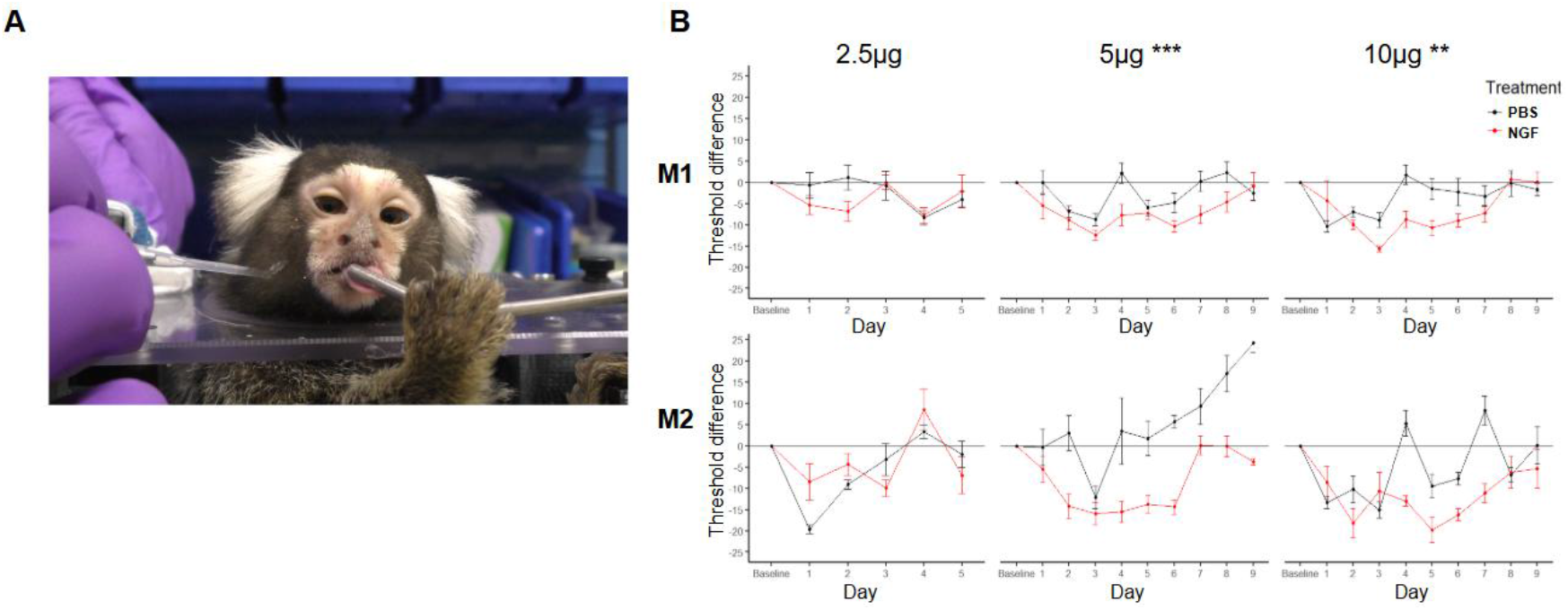
Mechanical withdrawal testing. (A) Example of marmoset seated in the primate chair with access to acacia gum while manually applying EVF to the masseter muscle. (B) Difference in mechanical withdrawal thresholds normalized to baseline values across post-injection behavioural testing (2.5 μg, 5 μg, 10 μg) on the NGF-injected (NGF, red) and saline-injected (PBS, black) masseter muscle for each monkey (M1, female; M2, male). Data presented as the mean ± standard error between treatments with ** p<0.01, and *** p<0.0001.

###### Maximum gape

Maximum gape was evaluated by offering a large high-reward food item and measuring the size of the animal’s bite. Animals were offered a jumbo marshmallow (cylindrical, 5 cm diameter, 4 cm height) while seated in the primate chair and aligned with a 12 cm ruler (Figure 3A). This alignment created a lateral view of each bite to be measured through video analysis. The maximum vertical gape of each bite was measured using video analysis, Video to Photo software (v.1.1.1.0), and ImageJ (42). Images were calibrated using a ruler attached to the marmoset chair and assessed frame-by-frame to determine the largest mouth opening for each bite attempt. The largest bite from each day was recorded as maximum gape and was normalized to an average baseline value from six days.

**Figure 3.**
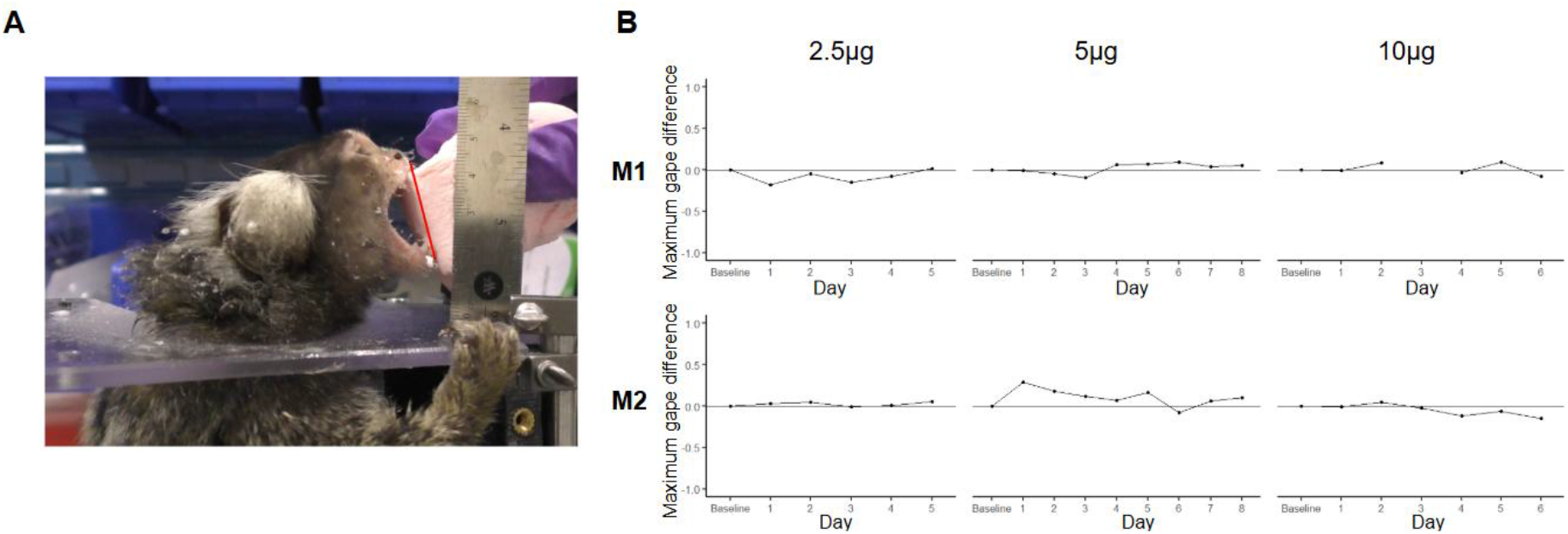
Maximum gape evaluation. (A) Example of a marmoset seated in the marmoset chair in the lab during gape measurements with a red line representing the distance measured through ImageJ. (B) Difference in the maximum gape normalized to baseline average across post-injection conditions (2.5 μg, 5 μg, 10 μg) for each monkey (M1, female; M2, male). Data presented as the normalized maximum value for each day with no significant differences between conditions.

###### Eating behaviour

Eating behaviour was assessed through a novel in-lab chewing assay and through monitoring the total number of food items eaten in the cage. For the chewing assay, five food items of varying hardnesses were offered to the animals twice (10 items total) while seated in the primate chair. Food preferences were established for the chewing assay by presenting various foods of varying hardnesses across many sessions. Animals were offered high value reward foods which were alternated to maintain novelty. If a food item was consistently rejected, it was excluded from the assay. Chewing was analyzed through video analysis following behavioural testing. Using BORIS (Behavioural Observational Research Interaction Software, v.8.27.7), food preference was determined by scoring if the animal chewed (exerted enough bite force to puncture the item) or did not chew each item. Items were also categorized and scored by their hardness or chewing difficulty from 1 to 5 (1: soft, easy to chew to 5: hard/chewy, difficult to chew) and each session had 1 food item from each ranking (Supplementary table 2). These data were visualized through the ggplot2 package in R to determine if animals changed their food preference to softer foods during treatment conditions. Additionally, animal care records were used to record the number of food items left in the cage for each day of testing across all baseline and experimental trials.

###### Grimace analysis

Grimace was assessed using DeepLabCut (v2.2.0) (43,44) to track the changes in ear tuft area and eye distance during each session. We manually labelled the corners of the ear tufts and eyelids on 50 frames from 11 videos (550 total) and trained a ResNet-50 neural network for 150,000 iterations (test error = 8.82, train error = 5.06, p-cutoff = 0.6) (Supplementary Video 1). Using RStudio, extracted coordinates (x,y) were converted into distances (cm) and each eye distance and tuft area was calculated across frames with any point with a likelihood below 0.90 removed as an outlier. Grimace was assessed across the testing session as well as during periods of withdrawals from mechanical threshold testing. Individual withdrawals were manually extracted and grouped according to whether the withdrawal occurred from palpation on the PBS-injected side, NGF-injected side, or neither (baseline).

Withdrawals were then averaged across each grouping.

### Experiment 2: Home cage behaviour analysis

#### NGF administration

Protocols developed during Experiment 1 were used for anesthesia, NGF reconstitution and administration. Based on the results of Experiment 1, 100 μl of NGF solution (100 μg/mL) or 100 μl of PBS was injected into the left or right masseter on Day 1 and Day 3.

#### Behavioural testing

Naturalistic home cage behaviours (circadian activity and spontaneous eating behaviour) were assessed prior to NGF treatment (before), during treatment (during) and after treatment (after). During this time, animals stayed in their home cage except for NGF injections (Day 1 and Day 3), sham anesthesia during baseline (Day 3), and mechanical withdrawal tests (Day 4) (Figure 1B). The duration of the NGF-treatment period was established using mechanical withdrawal thresholds from the 10 μg injection of Experiment 1 and the final day was considered as the final day that mechanical withdrawal thresholds were significantly lower on the NGF-injected muscle according to a linear mixed model (Day 7).

##### Home cage eating behaviour

Home cage behaviour recordings were performed during the morning feed for 30 minutes after the animals were presented with their food. Animals were housed in cages with a clear plexi-glass door and a video camera was manually placed daily to capture the eating shelf and approximately ¼ of the cage. Movement into the marmoset room was not restricted and marmosets were exposed to the standard daily activity of the animal care staff. Animal care records were used to record the number of food items left in the cage for each day of testing.

##### Circadian activity

Actiwatch^®^-Mini (CamTech, Cambridgeshire, UK, 128 kb, 24×7.7 mm, 7.5 g) was used to assess circadian rhythm by monitoring marmoset activity through uni-axial motion recordings (±0.05 g) collected every 10 seconds over a 24 h period for each animal (Figure 4B). They were placed on the neck of awake and restrained marmosets one day prior to the start of data collection and remained in place until the end of the recording period, except during anesthesia. Daytime rest, nighttime rest and activity metrics were downloaded and extracted using the SleepWare7 software. Daytime rest was defined as immobility for over 5 consecutive minutes during daytime hours (7:00-19:00) while nighttime rest was defined as immobility for over 5 consecutive minutes during nighttime hours (19:00-7:00).

**Figure 4.**
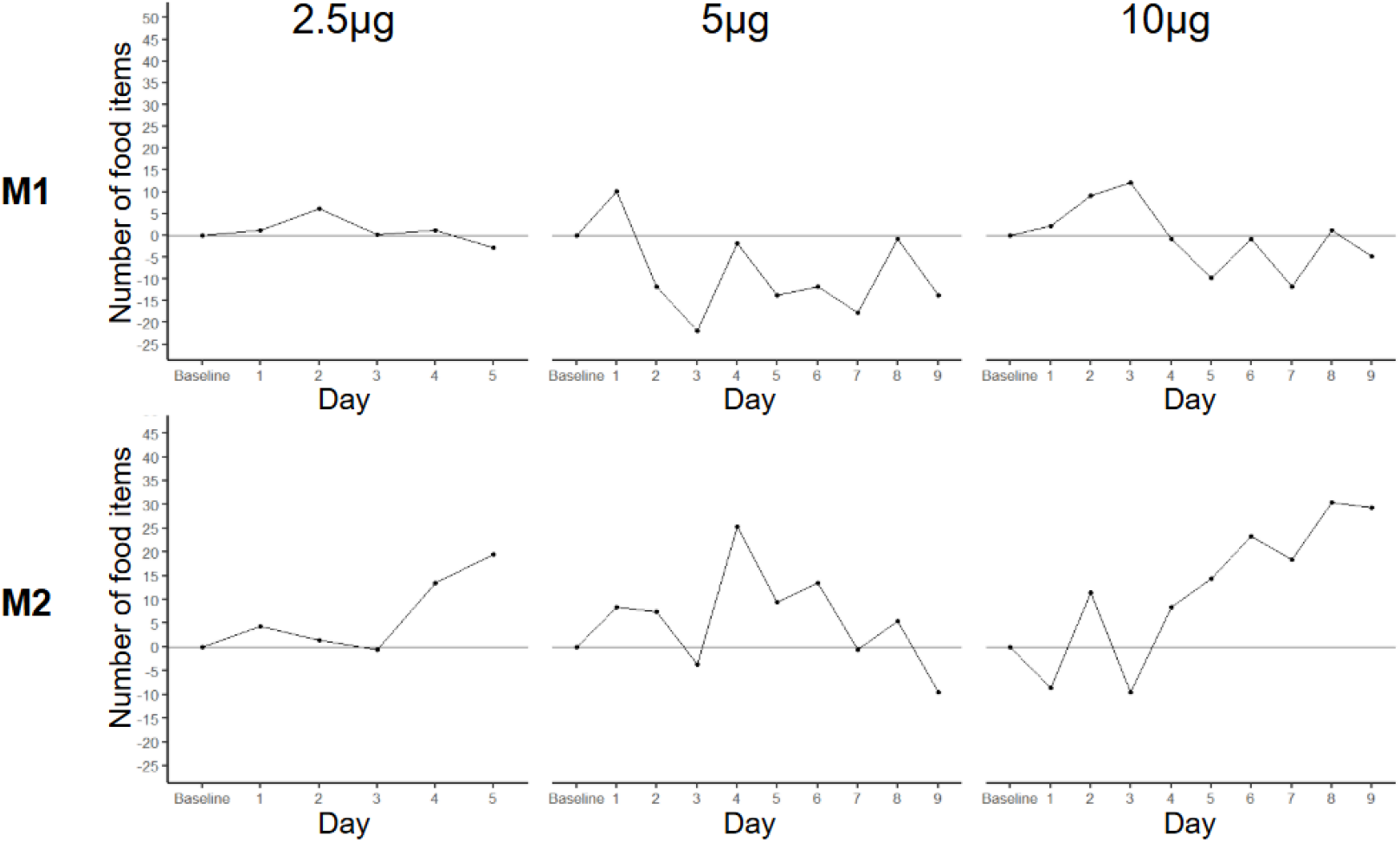
Difference in the number of food items left in the cage normalized to baseline values across each Experiment 1 post-injection condition (2.5 μg, 5 μg, 10 μg) for each monkey (M1, female; M2, male). Data presented as the normalized number of food items for each day with no significant differences between conditions.

#### Statistical analyses

All statistical analyses and graphs were generated using RStudio (v.4.3.1) (45).

For the mechanical withdrawal test, a linear mixed model with a random effect of animal, fixed effect of condition (baseline, 2.5 μg, 5 μg, 10 μg) and post hoc pairwise comparisons using estimated marginal means were used to compare normalized withdrawal thresholds between the saline-injected and NGF-injected muscle for each dose.

For the maximum gape evaluation, a linear mixed model with a random effect of animal, fixed effect of condition (baseline, 2.5 μg, 5 μg, 10 μg) and post hoc pairwise comparisons using estimated marginal means were used to compare maximum gape between baseline and each dose.

Eating behaviour was analyzed by comparing the number of food items left in the home cage during the morning feed. In Experiment 1, the number of food items not eaten were normalized to the average number of food items left in the cage during nine baseline in-lab testing days. In Experiment 2, the number of food items not eaten were normalized to the average number of food items left in the cage during thirty-one baseline days where animals were not removed from the cage. A generalized linear mixed model with a Poisson distribution, random effect of monkey, fixed effect of treatment, and post hoc pairwise comparison using estimated marginal means was used to compare the contrasts between the number of food items eaten during each dose and the respective baseline number of food items.

Grimace was analyzed using a linear mixed model with a random effect of animal, fixed effect of condition (baseline, 2.5 μg, 5 μg, 10 μg), and post hoc pairwise comparison using estimated marginal means to determine if there was a significant change in either the left or right ear tuft volume or eye distance across doses for the entire testing session or withdrawal periods only.

In Experiment 2, eating behaviours were analyzed through video analysis and manual behavioural scoring (BORIS; Supplementary table 3) (46). We used a linear mixed model with a fixed effect of condition (before, during, after), random effect of animal and post hoc pairwise comparisons using estimated marginal means for each of the following metrics, individually: time to first approach, time to first eating attempt, time to first eating instance, and percentage of food items dropped. We used a generalized linear mixed model using a Poisson distribution, random effect of animal, fixed effect of treatment, and post hoc pairwise comparisons using estimated marginal means for each of the following metrics, individually: number of total approaches and number of eating attempts.

Circadian activity was analyzed using the nparACT package (47) to assess nonparametric circadian rhythm analysis (NCPRA) including interdaily stability (IS), intradaily variability (IV), relative amplitude (RA), the average amplitude of the 5 hours with the lowest activity (L5), and the average amplitude for the 10 hours with the highest activity (L10) during each period for each animal. Additionally, we used a linear mixed model with a fixed effect of condition (before, during, after), random effect of animal and post hoc pairwise comparisons using estimated marginal means for each of the following extrapolated metrics, individually: total daytime rest time, average length of daytime rests, time of first daytime rest, time of first daytime rest after eating, average daytime activity, average activity 30 minutes before and 30 minutes after eating, and percentage of time spent resting during night time hours. We used a generalized linear mixed model using a Poisson distribution, random effect of animal, fixed effect of treatment, and post hoc pairwise comparisons using estimated marginal means for the following extrapolated metrics, individually: number of nighttime rest/awake bouts, number of daytime rests.

## Results

### Experiment 1: Dose effects and duration

#### Mechanical withdrawal

After injection of 100 μl of NGF (25 μg/mL, 50 μg/mL or 100 μg/mL) or PBS (control) into the left or right masseter on Day 1 and Day 3, relative mechanical withdrawal thresholds (Figure 2A) were compared between sides for each dose. A linear mixed model with a random effect of monkey revealed a significant interaction between dose and treatment and showed significantly lower relative mechanical thresholds on the NGF-injected side in the 5 μg (*p* < 0.0001) and 10 μg conditions (*p* = 0.001) but not the 2.5 μg condition when compared to PBS injected sides (Figure 2B). Post hoc analyses found this effect was driven by Day 4 (*p* = 0.0027), Day 6 (*p* = 0.0163), and Day 8 (*p* = 0.0369) in the 5 μg condition and Day 4 (*p* = 0.0001) and Day 7 (*p* = 0.0032) in the 10 μg condition. That is, 5 µg and 10 µg doses of NGF can evoke hypersensitivity starting the following day of the second injection (Day 4) and lasting for 3-4 days (Day 7-8).

#### Maximum gape

Maximum gape did not significantly differ from baseline conditions for either animal or in any dose according to a linear mixed model (Figure 3).

#### Eating behaviour

Eating behaviour did not show significant differences following NGF injection. Animals did not decrease the number of food items chewed in the lab and did not prefer softer foods when visually compared to baseline. Animals also did not show significant differences in the number of food items left in the cage in any treatment when compared with baseline conditions that included in lab testing (Figure 4). During both conditions, animals did not demonstrate any weight loss or sustained changes to observable behaviour (bright/active/responsive, quiet/active/responsive, or unresponsive).

#### Grimace

Grimace was not significantly different in any condition for either animal. Animals did not show significant changes in the left and right ear tuft area or eye distance across the entire testing session or across withdrawal periods (Supplementary Figure 1).

#### Experiment 2: Home cage behaviour analysis

Based on the results from Experiment 1, we injected 10 μg of NGF to further explore spontaneous behavioural changes in the home cage.

##### Home cage eating behaviours

Behavioural scoring of home cage eating behaviours revealed animals significantly increased the amount of time taken to eat the first food item during the treatment period (*p*_*Before-During*_ = 0.013, *p*_*During-After*_ > 0.038) (Figure 5A,B). Otherwise, animals did not significantly change their approach or eating behaviour during the treatment period for any of the other described behaviours. The number of food items left in the cage did not significantly increase during the treatment period (Figure 5C). However, the number of food items left in the cage was significantly affected by whether the animals were tested on in-lab assays during the testing period. Specifically, further analyses revealed that the number of food items left in the cage in Experiment 1 (which included daily in-lab testing) was significantly higher than the number of food items left in the cage during Experiment 2 (which did not include daily in-lab testing) (*p*_*10μg_without_inlab - 10μg_with_inlab*_ < 0.0001, *p*_*Baseline_without_inlab - Baseline_with_inlab*_ < 0.0001) (Figure 5D).

**Figure 5.**
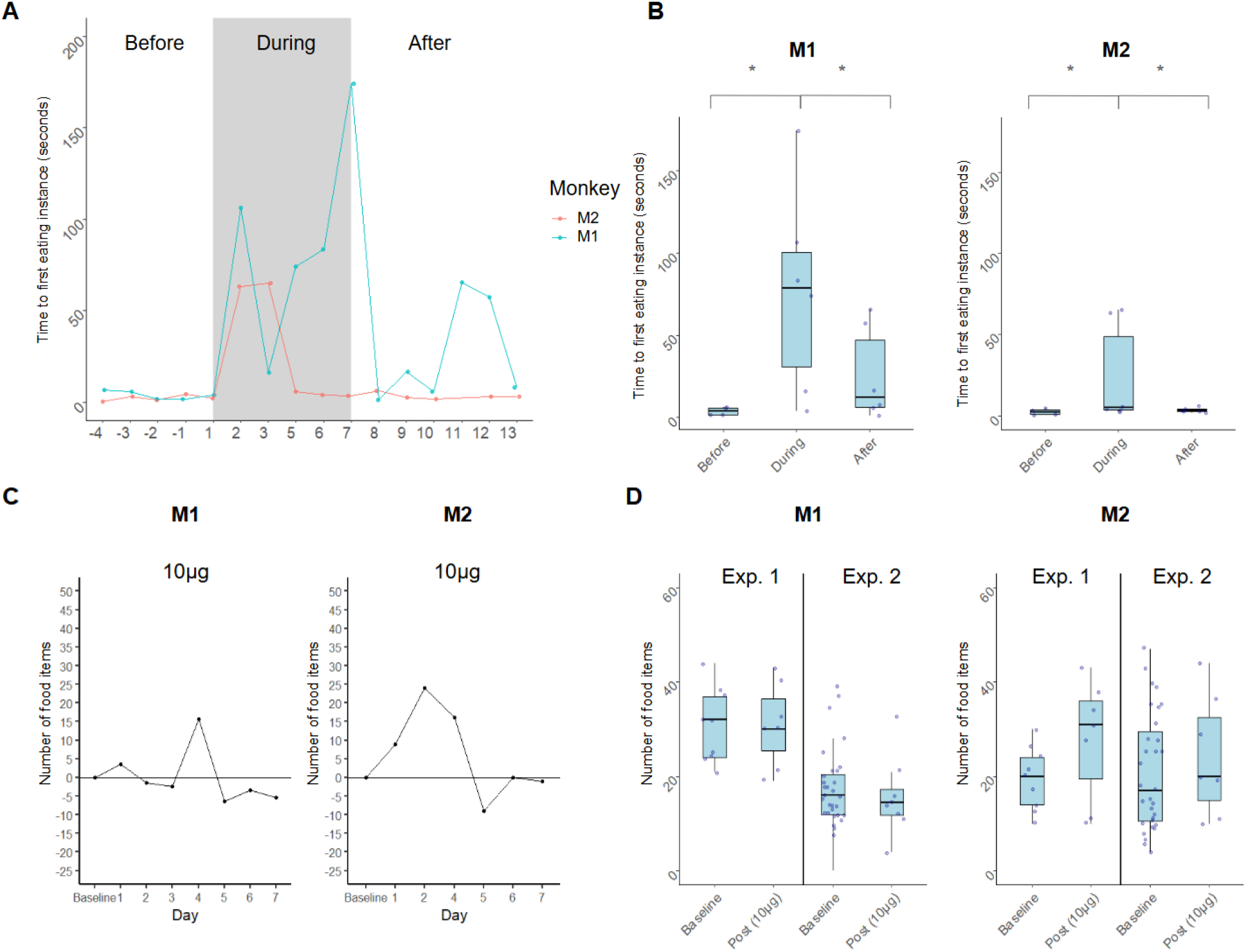
Home cage eating behaviour through video analysis for M1 (female) and M2 (male). (A) Amount of time to first eat a food item following the first approach to the food bowl across days for each animal (M1, M2). (B) Time needed to first eat a food item following the first approach to the food bowl across conditions (Before, During, After NGF treatment), presented as mean ± SD for M1 and M2. (C) Difference in the number of food items left in the cage normalized to baseline values across Experiment 2 post-injection conditions (10 μg) for each monkey (M1, M2). (D) Number of food items left in the cage for Experiment 1 (including in-lab behavioural testing) and Experiment 2 (excluding in-lab behavioural testing). Data presented with grey bars representing the treatment period and excluding the days animals were removed from the cage for in-lab testing.

##### Activity data

Visual assessment of NCPRA analysis found both animals had a higher intradaily variability (IV) during NGF injections which represents greater daily variability and a more disrupted circadian rhythm (Table 1; Figure 6A). However, given the degrees of freedom this could not be statistically confirmed and cannot be considered statistically significant. Further analyses were performed on activity data (nighttime rest, daytime rest) before, during and after treatment. Following NGF injection, animals significantly increased the total amount of daytime rest (*p*_*Before-During*_ <0.001, *p*_*During-After*_ = 0.015; Figure 6B) and the total number of daytime rests (*p*_*Before-During*_ = 0.004, *p*_*During-After*_ = 0.005; Figure 6C). Visual inspection of the average amplitude of daily activity suggested both animals had a minor decrease in activity amplitude during treatment conditions, although this effect was not significant (Figure 6D). There were no meaningful significant differences in the average length of daytime rests, time of first daytime rest, time of first daytime rest after eating, average activity 30 minutes before and 30 minutes after eating. Similarly, nighttime analyses did not exhibit significant differences in percentage of time spent or the number of rest/awake bouts.

**Table 1.** Actiwatch data showing interdaily stability (IS), intradaily variability (IV), relative amplitude (RA), the average amplitude for the 5 hours with the lowest activity (L5), and the average amplitude for the 10 hours with the highest activity (L10) for each monkey (M1, female; M2, male) and each period (before treatment, during treatment, after treatment).

| Monkey | Period | IS | IV | RA | L5 | M10 |
| --- | --- | --- | --- | --- | --- | --- |
| M1 | Before | 0.8 | 0.57 | 0.99 | 0.56 | 111.89 |
|  | During | 0.71 | 0.8 | 0.99 | 0.7 | 98.35 |
|  | After | 0.82 | 0.74 | 0.99 | 0.61 | 97.78 |
| M2 | Before | 0.73 | 0.49 | 0.98 | 0.56 | 63.28 |
|  | During | 0.71 | 0.58 | 0.98 | 0.4 | 51.18 |
|  | After | 0.85 | 0.51 | 0.99 | 0.25 | 63.52 |

**Figure 6.**
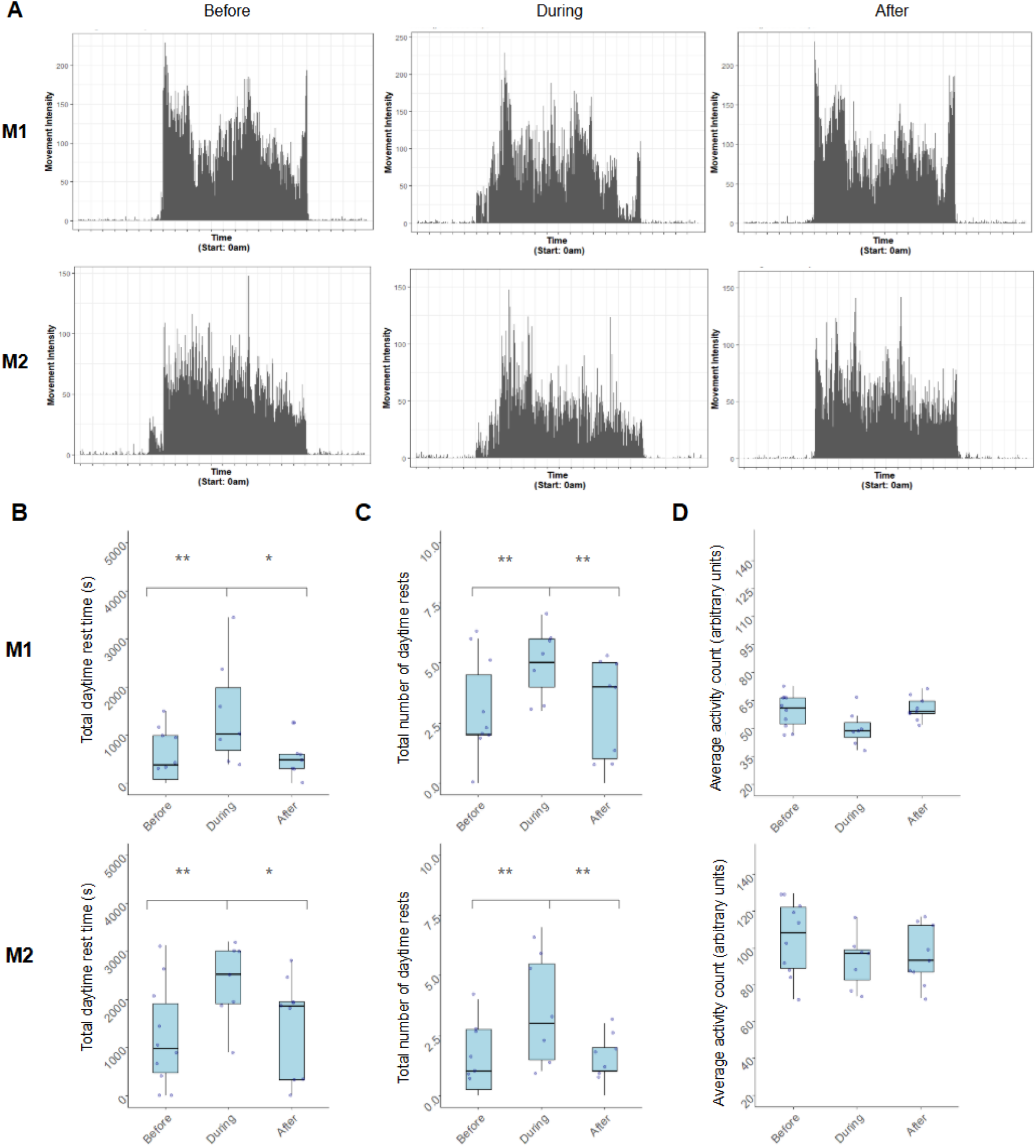
Activity data across conditions (before, during, and after treatment) for M1 (female) and M2 (male). (A) Average activity data across 24 hours (daytime hours: 7:00-19:00, nighttime hours: 19:007:00). (B) Total amount of daytime rest time. (C) Total number of daytime rests. (D) Average daily activity amplitude. Data presented as the significant difference between conditions with *p<0.05, and **p<0.01.

## Discussion

The goal of this study was to establish a model of myofascial TMD in the common marmoset using NGF and to assess subsequent pain behaviours. In Experiment 1, both animals showed changes in mechanical withdrawal thresholds on the NGF injected muscle after injection of 5 μg or 10 μg of NGF but did not show evidence of jaw dysfunction, altered chewing ability or grimace changes when tested on novel in-lab behavioural assays. However, in Experiment 2, both animals demonstrated differences in daily rest as well as minor changes to home cage eating behaviours after injection of 10 μg of NGF. The mechanical sensitivity we observed is characteristic of human and rodent NGF-TMD models (35,37–39,48). Additionally, considering we found minor changes in eating behaviour and circadian activity without changes to overall condition, weight or activity patterns, our model likely represents a similar model to the human which induces long-lasting mild pain.

In human and rodent studies, injection of NGF into the masseter induces allodynia or hyperalgesia through peripheral nerve sensitization (34,35,37,48). Mechanistically, NGF acts on tropomyosin kinase A receptors to lower the mechanical activation threshold of masticatory muscle afferent fibers (34). Injection of NGF into the masseter muscle increases the expression of peripheral NMDA receptors, which leads to an increase sensitivity of masticatory muscle afferent fibers to interstitial levels of the excitatory amino acid glutamate that may contribute to the prolonged mechanical sensitization produce by NGF (35,36). NGF has been shown to cause sustained reduced mechanical thresholds for up to two weeks following injection in rodent models (34,35,49) and increased pain ratings and hypersensitivity for up two weeks in human studies (37–40). Here, we validated this hypersensitivity in the marmoset by showing reduced mechanical withdrawal following injection of 5 μg or 10 μg of NGF. Both animals demonstrated lower pressure tolerance over the NGF injected muscle when compared to the saline injected muscle at higher NGF concentrations. Given mechanical hypersensitivity is a hallmark of myofascial TMD, this result demonstrates the efficacy of NGF and its ability to replicate this symptom in the marmoset. As such, our results validate the translation of this model into the common marmoset and provide an avenue to further explore possible cognitive, affective and behavioural pain measures.

Mechanical allodynia, however, is a single nociceptive component and does not account for other behavioural changes that can occur during the pain experience. Therefore, another goal of this study was to explore additional behavioural measures associated with the NGF-TMD model in the marmoset. Previous work in humans and rodents have observed that TMD related orofacial pain alters eating behaviours, jaw function and masticatory muscle pain sensitivity (11,12,37,48,50). Rodent studies have shown that orofacial pain can cause alterations in bite force (50), chewing time (12), and eating behaviours (such as eating duration and frequency) (11), while human studies have shown reduced gape and increased pain ratings during jaw function following injection of NGF (37–40). These masticatory effects correspond with decreases in von Frey hair withdrawal thresholds, therefore validating their use as behavioural markers for TMD related pain (11). Furthermore, the grimace scale has been widely used as a marker of pain across many animal species with recent work reducing subjectivity and labour costs through the integration of machine learning methods to track facial action units (51–57). Despite no validated grimace scale in the marmoset, previous studies have demonstrated ear position and orbital tightening as common markers of pain across mammal species (58) while marmoset facial action units have listed tuft upward, tuft downward and orbital tightening as action descriptors (59) and frequent facial movements (60). Based on these previous studies, in Experiment 1, we develop novel in-lab marmoset behavioural assays including a chewing assay to assess altered masticatory abilities, mouth opening task to assess for jaw dysfunction, and machine learning through DeepLabCut to track features of facial grimace. However, during treatment conditions, animals did not show reduced maximum gape, did not alter their chewing behaviour, and did not alter their ear tuft size or eye distance. These findings may be caused by the confounding stressors associated with laboratory testing masking spontaneous grimace changes and the high reward value of food items increasing motivation to bite food items despite orofacial pain. To address these limitations, in Experiment 2 we assessed behaviours in the home cage to explore naturalistic behaviours to regular diet food in a low stress environment. During the morning feed, animals increased the amount of time taken to eat the first food item but otherwise did not demonstrate other differences in approach behavior or the number of food items eaten daily under treatment conditions. This may suggest that while animals experience hesitancy to initiate feeding under treatment conditions, any discomfort or pain induced by NGF does not impact jaw functioning or masticatory abilities enough to alter their overall eating patterns in the home cage.

Another common comorbidity in chronic pain conditions — including in TMD (61) — is a disruption to sleep and circadian activity patterns. People with chronic pain often show a reciprocal relationship between sleep and pain disorders with pain leading to disrupted sleep patterns which amplifies the physiological, psychological and psychophysical aspects of the pain experience (62–66). This relationship in rodents describes changes to REM and NREM sleep duration, sleep fragmentation, sleep efficiency, and wakefulness in models of neuropathic pain (13,67), musculoskeletal pain (68), osteoarthritis (69), and orofacial pain induced by complete Freund’s adjuvant (CFA) (14). However, specific alterations to sleep or wakefulness patterns can vary between models, species, and subpopulations (13,67). In Experiment 2, we investigated possible interactions between NGF-induced pain and circadian activity in the marmoset, including with nighttime rest, daytime rest and daytime activity. We found both animals significantly increased the time spent resting during the day and the total number of daytime rests as well as showed evidence for a more fragmented daily rhythm. However, the number of nighttime rest bouts and the overall time spent resting during nighttime hours were not altered under treatment conditions. This may be due to the reduced severity of NGF pain, considering the human pain model demonstrates an average pain rating of 2.0/10 at rest (37), while the aforementioned studies use stimulants that generally report higher pain scores, such as CFA (14,70), hypertonic saline (68,71), or sciatic nerve crush (67,72). Additionally, previous studies often investigate sleep quality and patterns using electroencephalograms which can detect changes in REM or NREM sleep that cannot be quantified using accelerometry data.

Future studies should explore whether increased rest time during daytime hours during NGF-induced orofacial pain is associated with changes in REM or NREM sleep patterns rather than circadian activity.

### Limitations

The key limitation of this study is the small sample size (1 male, 1 female) which restricts the generalizability or interpretability of the findings. Past studies have demonstrated considerable sex differences in the pain response, specifically regarding the NGF-TMD model (37) which cannot be discerned through our findings. However, all the significant findings reported were found in both animals and future studies can address this limitation by increasing the number of both male and female marmosets included in the study. Another limitation is the effect of in-lab testing on naturalistic behaviours and subsequent behavioural assays. Specifically, we found significant differences in approach behaviours, home cage eating behaviours and the number of food items eaten in the home cage when animals performed in-lab testing, likely due to the stress of handling and food supplementation received during behavioural tasks. As such, any future studies should consider these confounding effects of in-lab behavioural assays when exploring naturalistic or spontaneous eating behaviours in marmoset models of orofacial pain.

Taken together, our results suggest that NGF can be used to induce hypersensitivity and yields minor changes to spontaneous behaviour in the marmoset. Although many of the behavioural assays did not show significant changes following NGF injection, the changes in circadian activity, eating initiation and hypersensitivity without debilitating impacts on overall condition, weight or general behaviour, support a pain model with a similar low pain rating as the human model. Our study is also one of the first to systematically investigate orofacial pain and related behaviours in the marmoset. We developed many novel behavioural assays in the marmoset, including machine learning methods of tracking marmoset grimace, and novel methods of investigating masticatory behaviours while also exposing the impact of in-lab testing on assessing spontaneous or minor exhibitions of pain. Overall, by investigating pain behaviours in the marmoset and determining a concentration and volume of NGF necessary to induce prolonged hypersensitivity, our findings establish an NGF-TMD non-human primate model similar to the human model which can facilitate future research on biomarkers (73) mechanisms, and interventions in TMD and related chronic pain disorders.

## Data availability

Data will be made available upon reasonable request.

## Supplementary material

Supplementary material: https://osf.io/jfz8q/?view_only=233a754682314abc8e91348df457232b

## Author contributions

**E.J.H., R.K., J.Y.K., B.E.C., J.A.P., D.A.S**. conceived and designed research; **E.J.H., R.K**. performed experiments; **E.J.H**. analyzed data; **E.J.H., R.K., M.B., J.A.P., D.A.S**. interpreted results of experiments, **E.J.H., M.B**. prepared figures; **E.J.H**. drafted manuscript; **E.J.H., R.K., M.B., J.Y.K., B.E.C., J.A.P., D.A.S**. edited and revised manuscript; **E.J.H., R.K., M.B., J.Y.K., B.E.C., J.A.P., D.A.S**. approved final version of manuscript.

