## Supplementary material for "Developing a model of temporomandibular disorder in the common marmoset using nerve growth factor"

**Table 1.** Concentrations and volumes used to establish an effective NGF dose.

| Injection Round | Concentration | Volume |
| --- | --- | --- |
| 1 | 25 µg/mL | 0.005mL |
| 2 | 25 µg/mL | 0.01mL |
| 3 | 25 µg/mL | 0.02mL |
| 4 | 25 µg/mL | 0.04mL |
| 5 | 25 µg/mL | 0.05mL |
| 6 | 25 µg/mL | 0.10mL |
| 7 | 50 µg/mL | 0.10mL |
| 8 | 100 µg/mL | 0.10mL |

**Table 2.** Food rating scale to assess in-lab food preferences from 1, representing food items that are the easiest to chew, to 5, representing items that are the most difficult to chew.

| Food Scale | Rating |
|---|---|
| Blackberry | 1 |
| Blueberry | 1 |
| Marshmallow | 1 |
| Banana | 1 |
| Pasta | 1 |
| Bean | 2 |
| Cantaloupe | 2 |
| Grapes | 2 |
| Mango | 2 |
| Pear | 2 |
| Apple | 3 |
| Caramel corn | 3 |
| Chicken | 3 |
| Cookie | 3 |
| Goldfish | 3 |
| Raisin | 3 |
| Fruit loop | 4 |
| Dried mango | 4 |
| Yogurt drop | 4 |
| Yogurt pretzel | 5 |
| Jellybean | 5 |
| Banana chip | 5 |
| Peanut brittle | 5 |

**Table 3.** Ethogram describing the eating behaviours assessed during home cage video analyses 30 minutes after the animal is presented with their morning feed.

| Behaviour | Measure | Type | Description |
|---|---|---|---|
| Approaches | First approach | Time | Amount of time to first approach the food bowl and engage with the contents (looking at the bowl and leaning inwards) following placement of the bowl in the cage. |
| | Approaches | Instance | Number of approaches to the food bowl defined by engagement with the contents (looking at the bowl and leaning inwards). |
| Eating attempts | First eating attempt | Time | Amount of time to first grab a food item with either the mouth or hand. |
| | Eating attempts | Instance | Number of times the animal grabs a food item either with the mouth or hand. |
| | Item dropped | Instance | Number of times the animal drops food item after item is grabbed |
| | Item eaten | Instance | Number of times the animal bites food item after item is grabbed |
| | | Time | Amount of time to first bite a food item after item is grabbed |
| | Item taken | Instance | Number of times the animal drops leaves the visible scene with item after item is grabbed |
| Marking | Marking | Instance | Number of times the animal marks as defined by an arched back and rubbing of the scent glands |

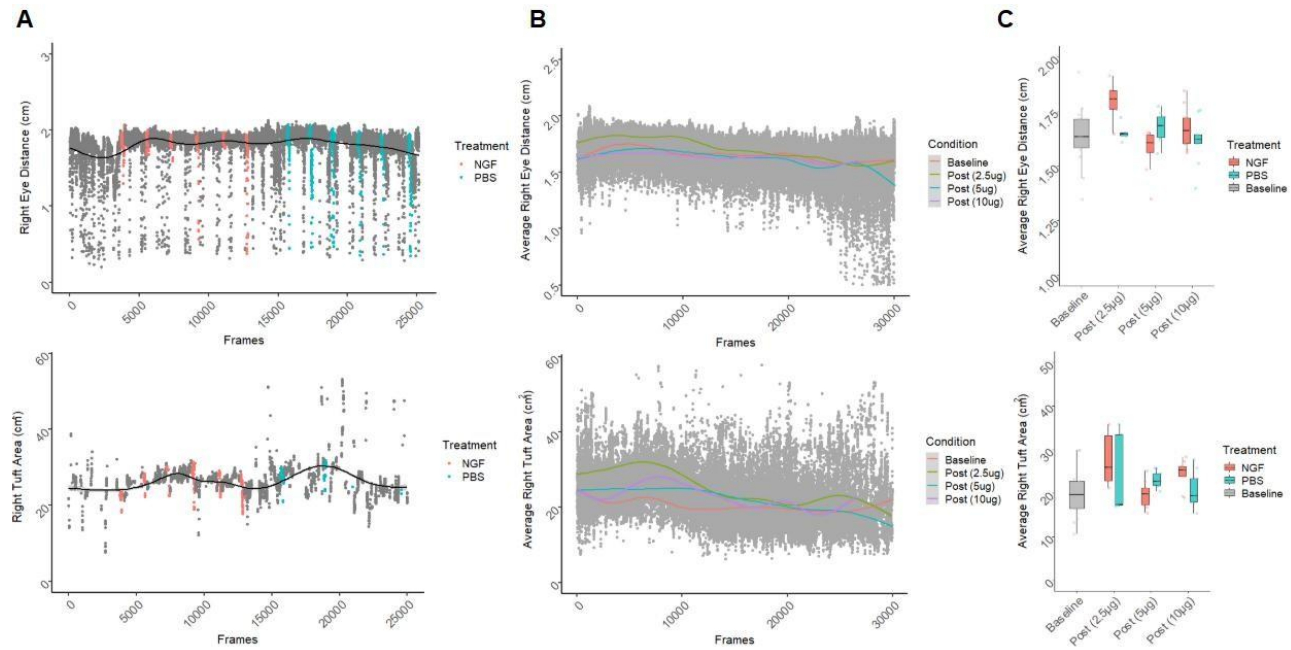


**Supplementary Figure 1.** Examples of grimace analyses for the right ear tuft and right eye distance on M1. (A) Right eye distance and right tuft area across frames of one randomly selected testing session with red and blue points representing the 2 second instances in which the animal withdrew during mechanical threshold testing on the NGF side or the PBS side. (B) Mean right eye distance or right tuft area across frames averaged across each dose (baseline, 2.5 µg, 5 µg, 10 µg) with lines representing a line of best fit (fitted using a general additive model). (C) Average right eye distance and right tuft area during only periods of withdrawals averaged across withdrawals during baseline (no treatment), on the PBS side, or on the NGF side (during treatment) for each dose (baseline, 2.5 µg, 5 µg, 10 µg).
